# Ancestry-dependent Enrichment of Deleterious Homozygotes in Runs of Homozygosity

**DOI:** 10.1101/382721

**Authors:** Zachary A. Szpiech, Angel C.Y. Mak, Marquitta J. White, Donglei Hu, Celeste Eng, Esteban G. Burchard, Ryan D. Hernandez

## Abstract

Runs of homozygosity (ROH) are important genomic features that manifest when an individual inherits two haplotypes that are identical-by-descent. Their length distributions are informative about population history, and their genomic locations are useful for mapping recessive loci contributing to both Mendelian and complex disease risk. We have previously shown that ROH, and especially long ROH that are likely the result of recent parental relatedness, are enriched for homozygous deleterious coding variation in a worldwide sample of outbred individuals. However, the distribution of ROH in admixed populations and their relationship to deleterious homozygous genotypes is understudied. Here we analyze whole genome sequencing data from 1,441 individuals from self-identified African American, Puerto Rican, and Mexican American populations. These populations are three-way admixed between European, African, and Native American ancestries and provide an opportunity to study the distribution of deleterious alleles partitioned by local ancestry and ROH. We re-capitulate previous findings that long ROH are enriched for deleterious variation genome-wide. We then partition by local ancestry and show that deleterious homozygotes arise at a higher rate when ROH overlap African ancestry segments than when they overlap European or Native American ancestry segments of the genome. These results suggest that, while ROH on any haplotype background are associated with an inflation of deleterious homozygous variation, African haplotype backgrounds may play a particularly important role in the genetic architecture of complex diseases for admixed individuals, highlighting the need for further study of these populations.

## Introduction

Runs of Homozygosity (ROH) are long stretches of identical-by-descent (IBD) haplotypes that manifest in individual genomes as the result of recent parental relatedness. Originally conceived to improve the accuracy of homozygosity mapping of recessive Mendelian diseases, ROH have formed the foundation of studies investigating the contribution of recessive deleterious variants to the genetic risk for complex diseases and the to the determination of complex traits [1]. Moreover, they have provided unique insights into the demographic and sociocultural processes [1] that have shaped genomic variation patterns in contemporary worldwide human populations [2–12], ancient hominins [13–16], non-human primates [17, 18], woolly mammoths [19], livestock [20–26], birds [27, 28], felines [29], and canids [30–39]. Recent population bottlenecks, cultural preferences for endogamy or consanguineous marriage, and natural selection, can create increased rates of ROH in individual genomes, substantially increasing overall homozygosity in such populations.

Several studies of the distribution of ROH in ostensibly outbred human populations have shown that ROH are common and range in size from tens of kilobases to several megabases in length [2–8]. Furthermore, total length and prevalence of ROH are correlated with distance from Africa [5, 7, 8], with more and longer ROH manifesting in individuals from populations a longer distance away. These patterns likely reflect increased IBD among haplotypes as a result of the serial bottlenecking process that humans experienced as they migrated out of Africa.

The prevalence of ROH in individual genomes has also been an important factor for understanding the genetic basis of complex phenotypes [40–43]. High levels of ROH have been associated with heart disease [44–47], cancer [44, 48-52], blood pressure [53–57], LDL cholesterol [57], various mental disorders [58–63], human height [64, 65], and increased susceptibility to infectious diseases [66]. Indeed, these results are consistent with the idea that many rare alleles of small effect may be the cause of increased risk for complex diseases [67–71], especially if these mutations are recessive [4, 72].

We have previously shown that ROH, especially long ROH, are enriched for deleterious homozygous variation [73, 74]. Whereas an overall increase in homozygotes is expected with increasing genomic ROH, we have shown that the rate at which deleterious homozygotes accumulate outpaces the rate at which benign homozygotes accumulate [73, 74] in long ROH (ROH on the order of several megabases). This result is a consequence of young (long) haplotypes with low-frequency variants segregating on them being paired IBD [74]. As low-frequency variants are more likely to be deleterious, the processes that create very long ROH can also generate unusually high numbers of deleterious homozygotes within these regions.

Although a few studies describing the worldwide distribution of ROH patterns have included a small number of admixed populations [5, 7, 8], the number of individuals per admixed population has been fairly small. Even as the number of admixed individuals continues to grow in the United States [75], they are still relatively understudied, which translates to disparities in our understanding of population-specific genetic factors that may influence complex phenotypes [76, 77]. Indeed, admixed populations have unique features compared to other populations, in that genomes from these populations are recent combinations of two or more ancestral populations.

This ancestral mosaicism has been exploited to make inferences about the natural history of human populations [78–88] and to search for ancestral haplotypes that influence complex phenotypes [89–96]. Here we add to the body of work on admixed populations by examining the relationship between ROH, local ancestry, and the accumulation of deleterious alleles. We use 1,441 recently published [97] whole genome sequences (dbGaP accession numbers phs000920 and phs000921) distributed roughly equally across three admixed populations in the Americas: African American (*n* = 475), Mexican American (*n* = 483), and Puerto Rican (n = 483). Each of these populations is three-way admixed between European, Native American, and African ancestral populations, although each has a distinct history.

Among the ancestral populations that contributed haplotypes to these admixed populations, it has been shown that the distribution of deleterious heterozygotes and deleterious homozygotes changes with distance from Africa [98–101]. With this in mind, we propose that accumulation of deleterious homozygotes via increased genomic ROH may also differ within admixed populations based on differing ancestral haplotypes. Indeed, with high deleterious heterozygosity, we propose that African ancestral haplotypes may be most susceptible to large increases in deleterious homozygotes when subjected to harsh bottlenecks or inbreeding, as these low frequency deleterious alleles will be paired into homozygotes as a result of increased genomic ROH.

## Results

### Admixture

Using the subset of sites from our whole-genome sequencing data that intersected with our African, European, and Native American reference panels, we called 3-way local ancestry tracts in all 1,441 samples (see Methods). We also estimated global ancestry proportions by summing the length of all haplotypes inferred to be from a given ancestry and dividing by the total genome length. Fig 1 summarizes the global ancestry proportions for all individuals from each population on a ternary plot. The admixture proportions largely accord with previous results in these populations, with Puerto Ricans having mostly African and European ancestry, Mexican Americans having mostly European and Native American ancestry, and African Americans having mostly African and European ancestry to the near exclusion of any Native American ancestry. However, although African Americans are frequently treated as a 2-way admixed population between European and African sources, we show that several AA individuals have non-trivial proportions of Native American ancestry. This suggests that, in general, a 2-way admixture model may not be uniformly appropriate for studying admixture patterns amongst self-identified African American individuals.

**Fig 1.**
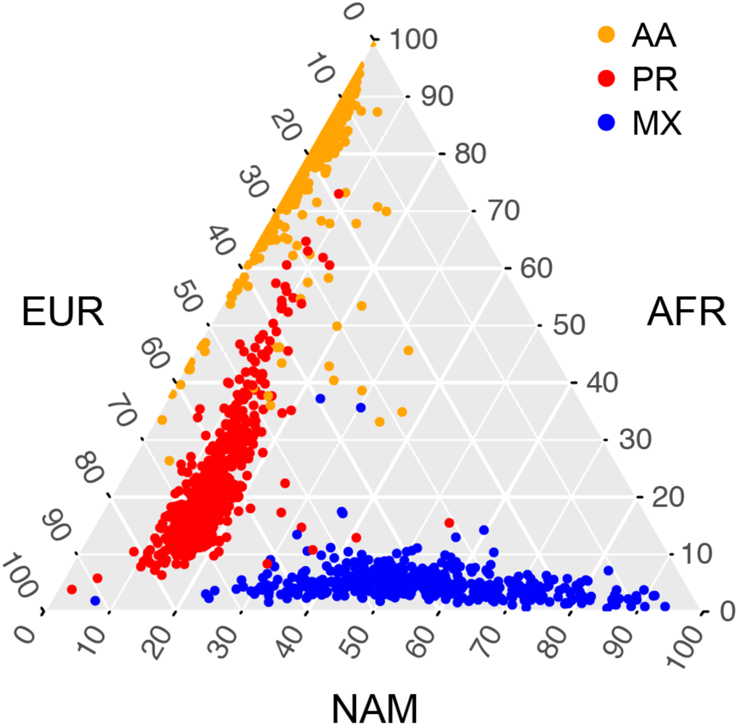
A ternary plot of global ancestry proportions. Each point represents a single individual, with their global ancestry proportions shown on each of the three axes (European, EUR; African, AFR; and Native American, NAM). Individuals are colored based on their reported ethnicity, with African Americans (AA) colored orange, Puerto Ricans (PR) colored red, and Mexican Americans (MX) colored blue.

### Runs of Homozygosity

We followed the ROH calling pipeline of Pemberton et al. [7] as implemented in the software GARLIC [102] to call ROH from the full whole-genome sequencing data (see Methods). This method identifies three classes of ROH based on the length distribution in each population. We refer to these size classes as short, medium, and long. These classes roughly correspond to ROH formed of IBD haplotypes from different time periods from the population history. Short ROH are tens of kilobases in length and likely reflect the homozygosity of old haplotypes; medium ROH are hundreds of kilobases in length and likely reflect background relatedness in the population; and long ROH are hundreds of kilobases to several megabases in length and are likely the result of recent parental relatedness. Total length of ROH in the genome is correlated with distance from Africa [4, 7]. In the case of our admixed populations, we therefore expect the total length of ROH to be correlated with increased European and Native American admixture fraction. Indeed, Fig 2A illustrates this pattern, with AA individuals having lowest total ROH, PR individuals having intermediate total ROH, and MX individuals having the highest total ROH (all pairwise Mann-Whitney U tests *p* < 2.2 × 10^−16^). Breaking down ROH by size class, we find that the total length of short ROH is comparable between PR and MX individuals (Fig 2B), but the total length of both medium ROH (Fig 2C) and total long ROH (Fig 2D) is highest on average in MX individuals.

**Fig 2.**
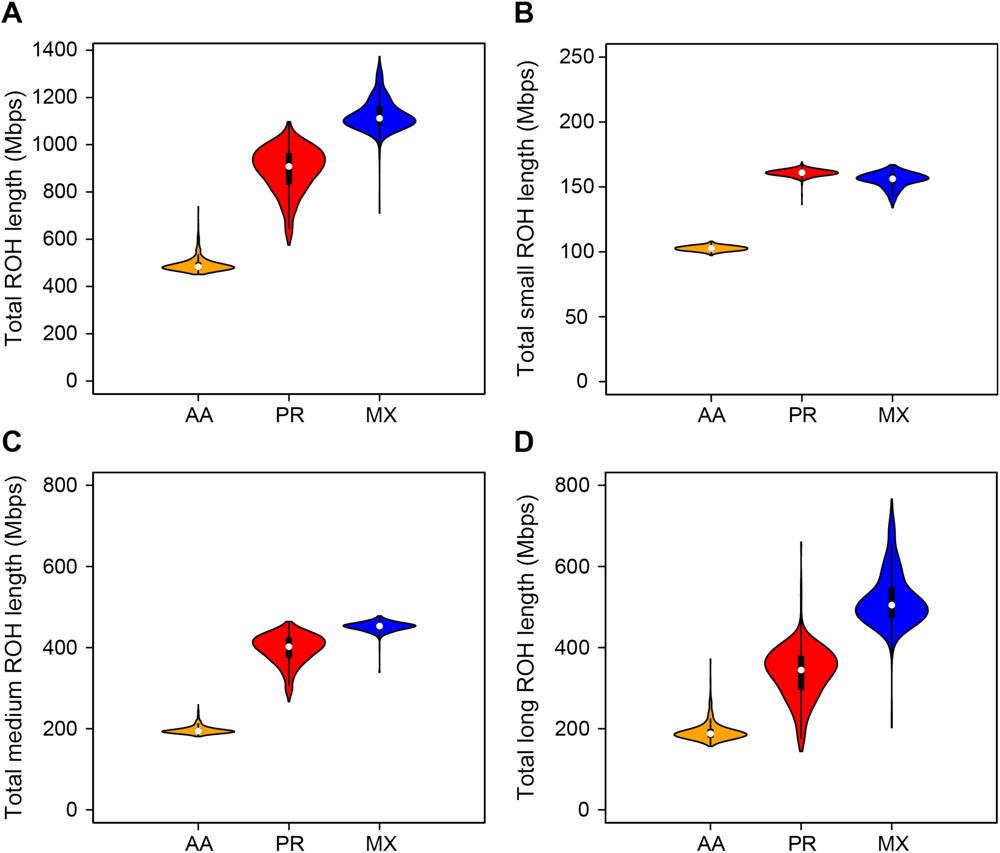
The distribution of summed ROH lengths across size classes. (A) all ROH, (B) short ROH, (C) medium ROH, and (D) long ROH. AA-African American, PR – Puerto Rican, MX – Mexican American.

### Deleterious Alleles

We used multiple approaches to predict the deleteriousness of all sites in the genome (see Methods), but focus on missense mutations classified as Probably Damaging, Possibly Damaging, or Benign using Polyphen 2 [103]. As in [73], we combine the Probably Damaging and Possibly Damaging mutations into a single “damaging” class, and we combine all Benign mutations with synonymous mutations into a single “benign” class. For individual *i* across all sites, we denote by 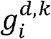 and 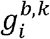 the total number of sites with *k* ∈ {0,1,2} alternate alleles classified as damaging or benign, respectively. In Fig 3A we plot the distribution of deleterious heterozygotes per individual, 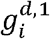, split by population. Consistent with previous work [98–101], we see an increased number of deleterious heterozygotes in populations with more African ancestry, with AA individuals having the most and MX individuals having the fewest (patterns replicate with other deleterious categories, see S5-S10 Figs). Conversely, we would expect an increase of deleterious homozygotes per individual in populations with more non-African ancestry. Indeed, in Fig 3B we plot the distribution of deleterious homozygotes per individual, 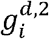, split by population and observe AA individuals with the fewest and MX individuals having the most (these patterns also replicate with other deleterious categories, see S5-S10 Figs). Figure 3C plots the total number of deleterious alleles per individual 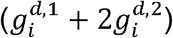. Contrary to other work [101], we find a total deleterious load highest on average in AA individuals. Although this pattern replicates across several other deleterious calling methods (S5-S9 Figs), when using GERP scores (as in [101]) the pattern reverses (S10 Fig) and is consistent with [101].

**Fig 3.**
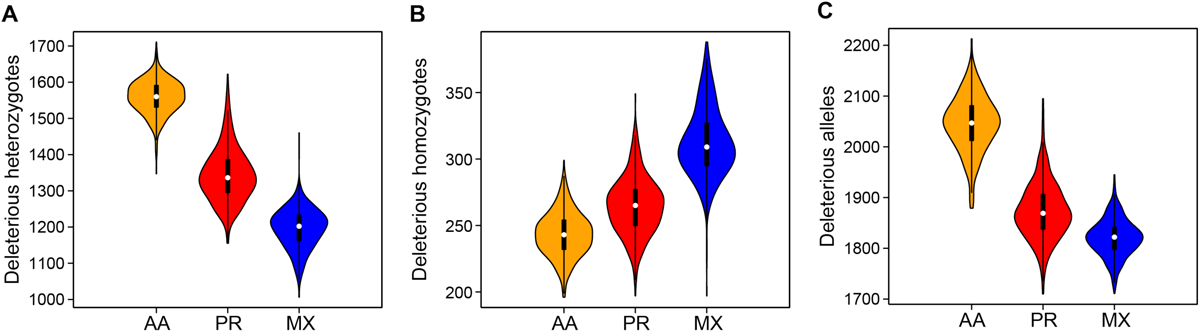
The distribution of deleterious alleles across populations. The number of (A) deleterious heterozygotes, (B) deleterious homozygotes, and (C) total deleterious alleles per individual using Polyphen2 classifications. AA – African American, PR – Puerto Rican, MX – Mexican American.

### Deleterious Alleles Across Local Ancestry

We next investigate whether there are any differences in deleterious load by local ancestry. Although our local ancestry calls provide us with phased local ancestry inferences, we were limited to a small subset of sites for our reference populations. Since the vast majority of our deleterious alleles come from our unphased whole-genome data, we do not have phase information for the deleterious alleles and cannot assign a specific ancestral haplotype in regions of discordant ancestry. Therefore, we calculate total load based on six different ancestry backgrounds. AFR, EUR, and NAM ancestry regions represent regions that are homozygous for African, European, and Native American ancestries, respectively, and AFEU, EUNA, and AFNA ancestry regions represent regions that are called heterozygous for African/European, European/Native American, and African/Native American ancestries, respectively. We then calculate for each population the number of deleterious alleles per basepair for each ancestry background.

Table 1 shows the number of deleterious alleles per basepair for each population and each ancestry background. We perform two types of tests for independence in order to determine whether there are significant differences in the number of deleterious alleles per basepair. First, we test for independence of the count of deleterious alleles on an ancestry background and the count of basepairs covered by that ancestry across populations. We find that neither African ancestry nor European ancestry have statistical differences in the number of deleterious alleles per MB across populations. Further, while NAM, EUAF, and AFNA exhibit statistically differences across populations, it appears to be driven by one of the two populations (AA, MX, and PR, respectively). Next, we test for independence of these counts across ancestries within each population. Here we find that all populations have statistically significant differences in the distribution of deleterious alleles across ancestry backgrounds (AA *p* < 2.2 x 10^−16^; MX *p* < 2.2 = 10^−16^; PR *p* < 2.2 × 10^−16^), with NAM ancestry having the lowest rate in AA and PR individuals and EUR having the lowest rate in MX individuals. However, we note that the overall differences were very small (a difference of < 0.1 deleterious alleles per Mbp).

**Table 1.** The number of deleterious alleles per megabase partitioned by population and local ancestry background.

| | AFR<br>($p = 0.160$) | EUR<br>($p = 0.452$) | NAM***<br>($p = 3.314 \times 10^{-7}$) | EUAF**<br>($p = 1.131 \times 10^{-3}$) | EUNA<br>($p = 0.123$) | AFNA**<br>($p = 4.392 \times 10^{-3}$) |
| --- | --- | --- | --- | --- | --- | --- |
| AA***<br>($p < 2 \times 10^{-16}$) | 0.335<br>($1.642 \times 10^6$) | 0.284<br>($1.009 \times 10^5$) | 0.237<br>($8.648 \times 10^2$) | 0.311<br>($7.943 \times 10^5$) | 0.280<br>($2.491 \times 10^4$) | 0.315<br>($8.364 \times 10^4$) |
| PR***<br>($p < 2 \times 10^{-16}$) | 0.337<br>($1.603 \times 10^5$) | 0.282<br>($1.064 \times 10^6$) | 0.275<br>($5.395 \times 10^4$) | 0.313<br>($7.517 \times 10^5$) | 0.286<br>($4.912 \times 10^5$) | 0.308<br>($1.700 \times 10^5$) |
| MX***<br>($p < 2 \times 10^{-16}$) | 0.341<br>($7.651 \times 10^3$) | 0.282<br>($4.585 \times 10^5$) | 0.286<br>($8.275 \times 10^5$) | 0.317<br>($1.154 \times 10^5$) | 0.287<br>($1.142 \times 10^6$) | 0.314<br>($1.393 \times 10^5$) |

Total number of megabases, summed across all individuals, in parentheses. A significant difference (Pearson's Chi-squared test, p-value in parentheses) across populations for a given ancestry background is denoted at the beginning of a column. A significant difference across ancestry backgrounds for a given population (Pearson's Chi-squared test, p-value in parentheses) is denoted at the beginning of a row. Population codes: AA - African American, PR - Puerto Rican, MX - Mexican American. Local ancestry codes: AFR - Homozygous African, EUR - Homozygous European, NAM - Homozygous Native American, EUAF - Heterozygous European/African, EUNA - Heterozygous European/Native American, AFNA - Heterozygous African/Native American. \* $p < 0.05$ , \*\* $p < 0.01$ , \*\*\* $p < 0.001$ .

### Deleterious Alleles in ROH

Next, we turn to examining the distribution of deleterious homozygotes within ROH. It was previously reported [73, 74] that there is a higher proportion of deleterious homozygotes per unit increase of ROH than expected from the proportion of benign homozygotes. Naturally, as the total amount of genomic ROH increases, we expect more homozygotes to fall within ROH. However, [73] and [74] found that the rate of increase of the proportion of deleterious homozygotes was greater than for benign homozygotes. This effect was strongest for long ROH, which are likely the result of recent parental relatedness.

For each individual *i* and for each ROH class *j* ∈ {*A, B, C, R, N*} (A - short ROH, B - medium ROH, C - long ROH, R - all ROH, and N - outside ROH), we define the number of damaging or benign sites with *k* ∈ {0,1,2} alternate alleles as 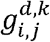 and 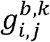, respectively. Thus we calculate the proportion of damaging homozygotes in ROH class *j* as

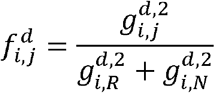

and the proportion of benign homozygotes in ROH as

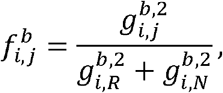

respectively. We also compute, for each individual *i* and each class *j*, the fraction of the genome covered in ROH as

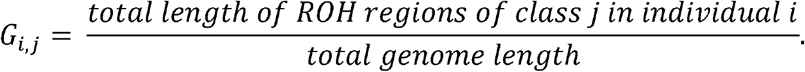

We plot the proportions of ROH homozygotes versus genomic fraction of ROH in Fig 4, which is analogous to Fig 4 from [73]. In order to determine if there is a statistically significant difference in the accumulation of deleterious homozygotes versus benign homozygotes, we construct a linear regression model (as in [73, 74]), *f.,_j_* = *β*_0_ + *β*_1_ *G.,_j_* + *β*_2_*D* + *β*_3_*DG.,_j_ + ε*, where *f.,_j_* is a vector of length 2,882 containing the proportions of both damaging and benign homozygotes in ROH class *j* for all individuals, *G.,_j_* is a vector of genomic class *j* ROH proportions, and *D* is an indicator variable taking a value of 1 when the response represents damaging homozygotes and 0 for benign homozygotes. In this framework, a statistically significant *β*_2_ suggests an overall higher proportion of damaging homozygotes in ROH compared to benign homozygotes, e.g. *β*_2_ = 0.1 means that an extra 10% of genome-wide deleterious homozygotes fall in ROH compared to the distribution of benign homozygotes. A statistically significant *β*_3_ suggests a difference in the rate of accumulation per unit increase of ROH, e.g. *β*_3_ = 1.0 means that for a 10% increase in genomic ROH, 10% more deleterious homozygotes fall in ROH compared to benign homozygotes. Inferred coefficients for the four regressions corresponding to *j* ∈ {*A, B, C, R*} each are given in Table 2.

**Table 2.** Regression coefficients inferred for the analyses shown in Fig 4 with p-values in parentheses.

| ROH<br>Class | $\beta_0$<br>(p-value) | $\beta_1$<br>(p-value) | $\beta_2$<br>(p-value) | $\beta_3$<br>(p-value) |
| --- | --- | --- | --- | --- |
| All | ***$3.122 \times 10^{-2}$<br>($< 2 \times 10^{-16}$) | ***1.460<br>($< 2 \times 10^{-16}$) | ***0.180<br>($< 2 \times 10^{-16}$) | $1.807 \times 10^{-2}$<br>(0.0671) |
| Long | $1.059 \times 10^{-3}$<br>(0.508) | ***1.429<br>($< 2 \times 10^{-16}$) | ***$5.335 \times 10^{-2}$<br>($< 2 \times 10^{-16}$) | ***0.229<br>($< 2 \times 10^{-16}$) |
| Medium | ***$8.510 \times 10^{-3}$<br>($< 3.86 \times 10^{-5}$) | ***1.584<br>($< 2 \times 10^{-16}$) | ***$7.012 \times 10^{-2}$<br>($< 2 \times 10^{-16}$) | *$5.695 \times 10^{-2}$<br>(0.0173) |
| Short | ***$1.265 \times 10^{-2}$<br>($4.61 \times 10^{-7}$) | ***1.424<br>($< 2 \times 10^{-16}$) | ***$4.810 \times 10^{-2}$<br>($< 2 \times 10^{-16}$) | ***-0.428<br>($1.10 \times 10^{-8}$) |

\* $p < 0.05$ , \*\* $p < 0.01$ , \*\*\* $p < 0.001$ .

**Fig 4.**
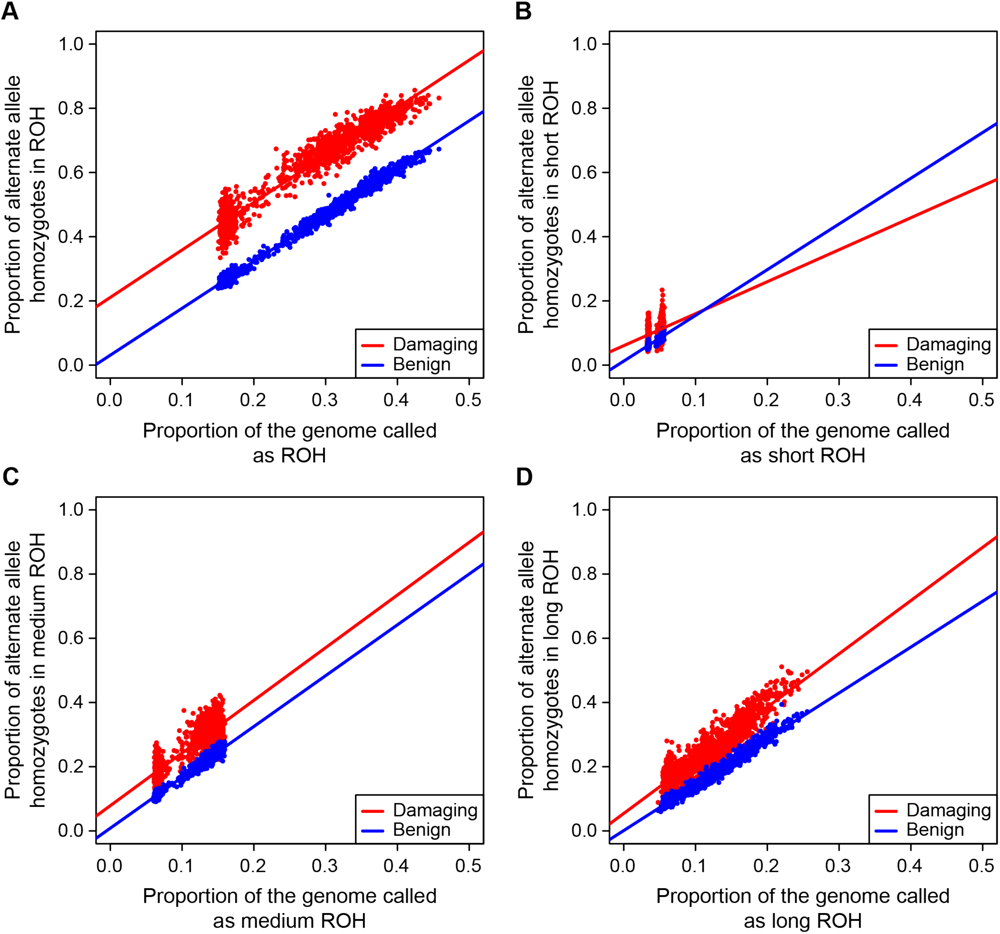
Deleterious and benign homozygotes in ROH classes. The proportion of damaging (red) and benign (blue) homozygotes falling in ROH of different size classes: (A) all ROH, (B) short ROH, (C) medium ROH, and (D) long ROH.

Fig 4A plots these proportions versus total ROH for all ROH classes combined. In agreement with [73], we find that there is an overall greater proportion of damaging homozygotes in ROH compared to benign homozygotes (*β*_2_ = 0.1799, *p* < 2 × 10^−16^), but in contrast the overall rate of accumulation is not different (*β*_3_ = 1.807 × 10^−2^, *p* = 0.0671). When we partition ROH by size class, the distribution of homozygotes in short ROH (Fig 4B) also differs from [73]. Whereas previously there were no statistically significant differences in *β*_2_ or *β*_3_, here we find a significant positive *β*_2_ = 4.810 × 10^−2^ (*p* < 2 × 10^−16^) and a statistically significant negative *β*_3_ = −0.428 (*p* = 1.10 × 10^−8^) suggesting that ROH comprised of old haplotypes accumulate deleterious homozygotes at a slower rate that benign homozygotes. As we expect short ROH to be comprised of old haplotypes that have been segregating for a long time, it is reasonable to think that only haplotypes with relatively few deleterious alleles remain segregating in the population. Our results for medium (Fig 4C) and long ROH (Fig 4D) are consistent with previous work [73, 74]; in particular we find that the difference in rates of gain of deleterious versus benign homozygotes is greatest in long ROH (*β*_3_ = 0.229; *p* < 2 × 10^−16^).

### Deleterious Alleles in ROH Partitioned by Local Ancestry

Now we turn to analyzing the distribution of deleterious homozygotes in ROH comprised of only one particular ancestral haplotypes. As shown in Fig 3A and in other work [98–101], populations with more African ancestry tend to have high numbers of deleterious heterozygotes genome-wide. This contrasts with populations that have more European and Native American ancestry, which tend to have more genome-wide deleterious homozygotes (Fig 3B) as a result of the serial bottlenecks they experienced since migrating out of Africa.

However, admixed populations are a recent combination of two or more ancestral populations, and since genome-wide ancestry proportions correlate with numbers of deleterious heterozygotes and homozygotes, we desire to investigate how this mosaicism might affect the accumulation of deleterious homozygotes in ROH. We have already shown (Fig 4) that as total genomic ROH increases the proportion of deleterious homozygotes falling in ROH increases faster than the proportion of benign homozygotes, but here we want to know if the ancestral background of the IBD haplotypes matters. Here we propose that haplotypes sourced from ancestral populations with high deleterious heterozygosity have highest rates of accumulation of deleterious homozygotes when paired IBD to generate ROH.

Why might we expect high deleterious heterozygosity haplotypes to generate large numbers of deleterious homozygotes in ROH? Pemberton and Szpiech [74] recently demonstrated that long ROH are enriched for homozygotes comprised of low-frequency alleles. Low-frequency alleles are more likely to be deleterious and are more likely to manifest in individual genomes as heterozygotes. Under a typical random mating scenario these low frequency alleles would be likely to segregate in the population largely as heterozygotes, however severe bottlenecks and cultural practices such as endogamy and consanguineous marriage substantially raise the likelihood of pairing low-frequency alleles as IBD homozygotes. We therefore expect that deleterious homozygotes will be concentrated in large proportion within ROH comprised of African ancestral haplotypes, and that the rate of gain of deleterious homozygotes will be greatest in ROH of African ancestral haplotypes.

To test this proposition, we first partition ROH based on the ancestral background of the underlying IBD haplotypes. Then we compute for each individual (*i*) the fraction of all deleterious (*d*) and benign (*b*) homozygotes across the genome that fall into each ROH class (*j*) as:

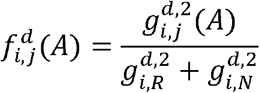

and

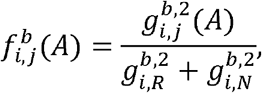

where 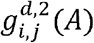 and 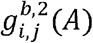 are the number of deleterious and benign homozygotes, respectively, in individual *i* in ROH class *j* on ancestral haplotype background *A* ∈ {*AFR,EUR,NAM*}. Similarly, 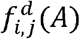 and 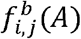 are the genome-wide fraction of deleterious and benign homozygotes, respectively, in individual *i* in ROH class *j* that fall on haplotype background A. Finally, we fit a linear model similar as above, *f.,_j_*(*A*) = *β*_0_ + *β*_1_*G.,_j_*(*A*) + *β*_2_*D* + *β*_3_*DG.,_j_*(*A*) + *ε*, in order to test for differences in the rate of accumulation (*β*_3_) of deleterious homozygotes compared to benign homozygotes as a function of *G.,_j_*(*A*), the genomic fraction of ROH on ancestral background A. The results are plotted in Fig 5 for total ROH (*j = N*; Fig 5A-C) and for long ROH (*j = C*; Fig 5D-F), and the regression coefficients are also summarized in Table 3.

**Table 3.**
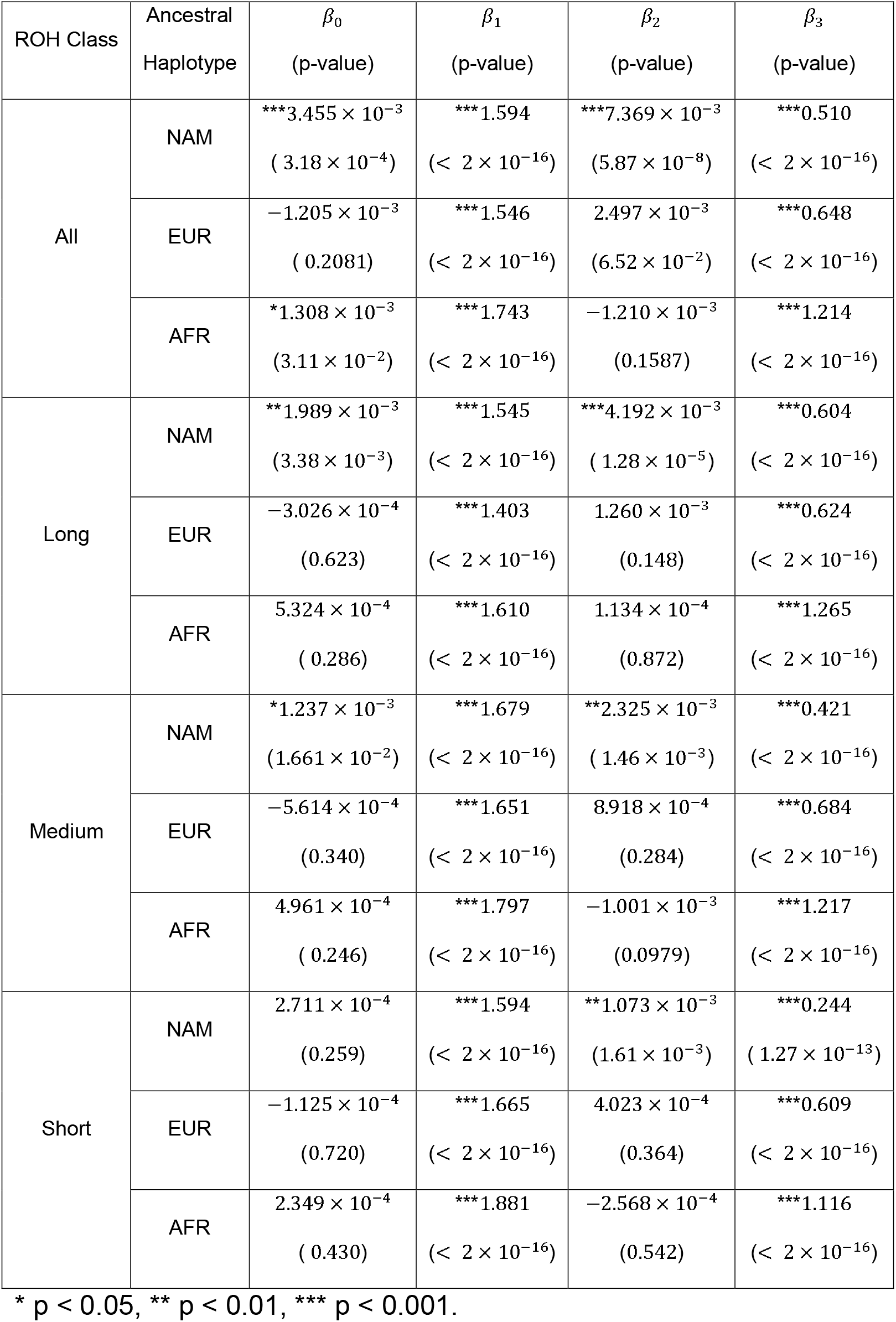
Regression coefficients inferred for the analyses shown in Fig 5 and S1 Fig with p-values in parentheses.

**Fig 5.**
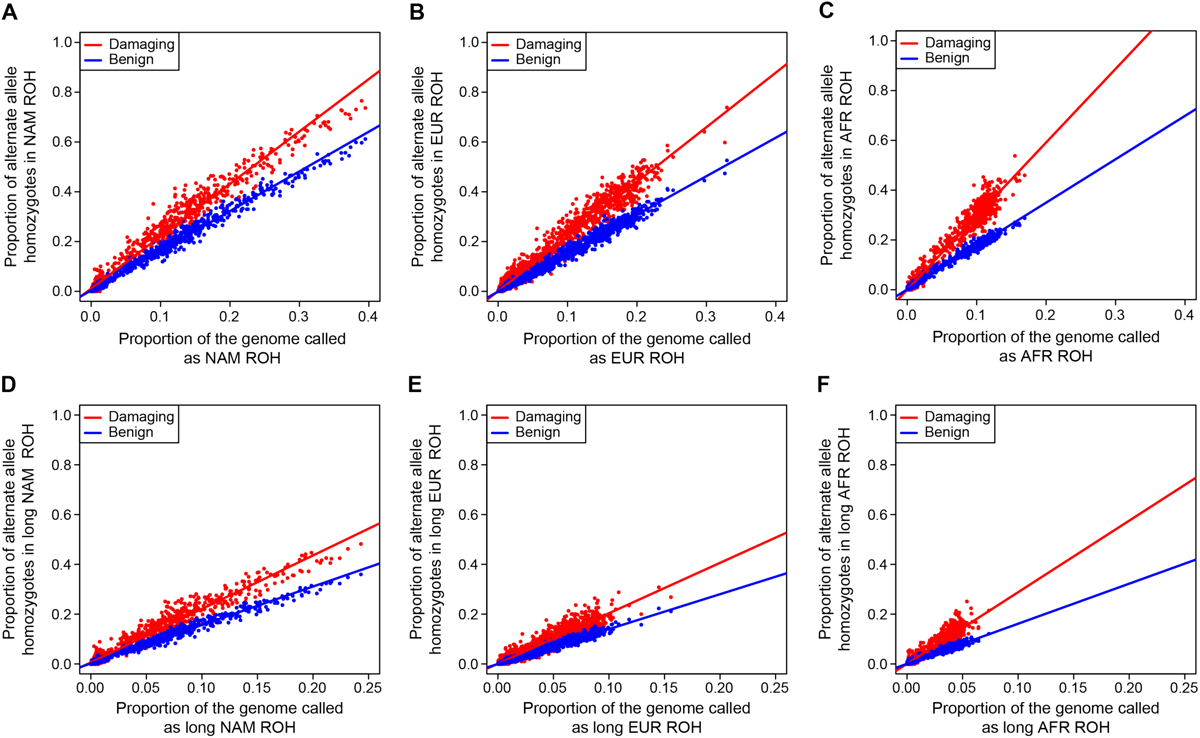
Deleterious and benign homozygotes in ROH classes separated by ancestry. The proportion of damaging (red) and benign (blue) homozygotes falling in ROH comprised of different ancestral haplotypes and size classes: (A) all NAM ROH, (B) all EUR ROH, (C) all AFR ROH, (D) long NAM ROH, (E) long EUR ROH, and (F) long AFR ROH. EUR – European, AFR – African, and NAM – Native American.

For total ROH, we find significant differences in the rate of accumulation of deleterious homozygotes on all ancestry backgrounds (Fig 5A-C). Furthermore, consistent with our expectations, we find that ROH on African ancestral haplotypes have the highest rate difference (*β*_3_ = 1.214, *p* < 2 × 10^−16^; Fig 5C), whereas ROH on European ancestral haplotypes have an intermediate rate difference (*β*_3_ = 0.648, *p* < 2 × 10^−16^; Fig 5B) and ROH on Native American ancestral haplotypes have the lowest rate difference (*β*_3_ = 0.510, *p* < 2 × 10^−16^; Fig 5A). This pattern is repeated when we consider only long ROH comprised of young haplotypes (Fig 5D-F) and also when we analyze smaller ROH (albeit with weaker effects; S1 Fig).

We next directly compare the rate of increase of deleterious homozygotes across different ancestral haplotype backgrounds. To do this we compute the following regression, 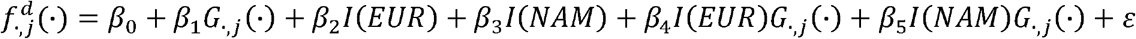, where 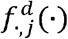 is a vector representing the proportion of damaging homozygotes in ROH class *j* on each local ancestry background across all individuals. *G.,_j_*(·) represents the genome-wide fraction ROH class *j* falling on each local ancestry background across all individuals, and *I*(*A*) is an indicator variable which takes the value 1 if the associated response is on ancestral background *A* ∈ {*AFR,EUR, NAM*} and takes the value 0 otherwise. Here we analyze each ROH class: all, long, medium, and short.

We plot the results for “all” and “long” in Fig 6 (“medium” and “short” in S2 Fig) and summarize the inferred regression coefficients for all classes in Table 4. We focus on the regression coefficients *β*_4_ and *β*_5_, which represent the difference in rate of gain of deleterious homozygotes in ROH on European or Native American haplotypes compared to African haplotypes, respectively. Graphically, in Fig 6 and S2 Fig, a significant *β*_4_ corresponds to a significant difference in the slope of the orange and blue line, and a significant *β*_5_ corresponds to a significant difference in the slope of the orange and purple line. Since we expect that the rate of gain of deleterious homozygotes to be lowest in ROH on European and Native American haplotypes compared to ROH on African ones, we expect significant negative values for both *β*_4_ and *β*_5_.

**Table 4.** Regression coefficients inferred for the analyses shown in Fig 6 and S2 Fig with p-values in parentheses.

| ROH Class | $\beta_0$<br>(p-value) | $\beta_1$<br>(p-value) | $\beta_2$<br>(p-value) | $\beta_3$<br>(p-value) | $\beta_4$<br>(p-value) | $\beta_5$<br>(p-value) |
| --- | --- | --- | --- | --- | --- | --- |
| All | $9.861 \times 10^{-5}$<br>(0.922) | ***2.957<br>($< 2 \times 10^{-16}$) | $1.193 \times 10^{-3}$<br>(0.431) | ***$-1.072 \times 10^{-2}$<br>($1.87 \times 10^{-12}$) | ***-0.763<br>($< 2 \times 10^{-16}$) | ***-0.852<br>($< 2 \times 10^{-16}$) |
| Long | $5.002 \times 10^{-4}$<br>(0.449) | ***2.878<br>($< 2 \times 10^{-16}$) | $4.477 \times 10^{-4}$<br>(0.661) | ***$5.453 \times 10^{-3}$<br>($7.11 \times 10^{-8}$) | ***-0.852<br>($< 2 \times 10^{-16}$) | ***-0.727<br>($< 2 \times 10^{-16}$) |
| Medium | $-4.804 \times 10^{-4}$<br>(0.443) | ***3.013<br>($< 2 \times 10^{-16}$) | $8.098 \times 10^{-4}$<br>(0.387) | ***$4.002 \times 10^{-3}$<br>($2.38 \times 10^{-5}$) | ***-0.678<br>($< 2 \times 10^{-16}$) | ***-0.912<br>($< 2 \times 10^{-16}$) |
| Short | $-2.196 \times 10^{-5}$<br>(0.953) | ***2.996<br>($< 2 \times 10^{-16}$) | $3.114 \times 10^{-4}$<br>(0.571) | *$1.351 \times 10^{-3}$<br>(0.0149) | ***-0.723<br>($< 2 \times 10^{-16}$) | ***-1.158<br>($< 2 \times 10^{-16}$) |

\* $p < 0.05$ , \*\* $p < 0.01$ , \*\*\* $p < 0.001$ .

**Fig 6.**
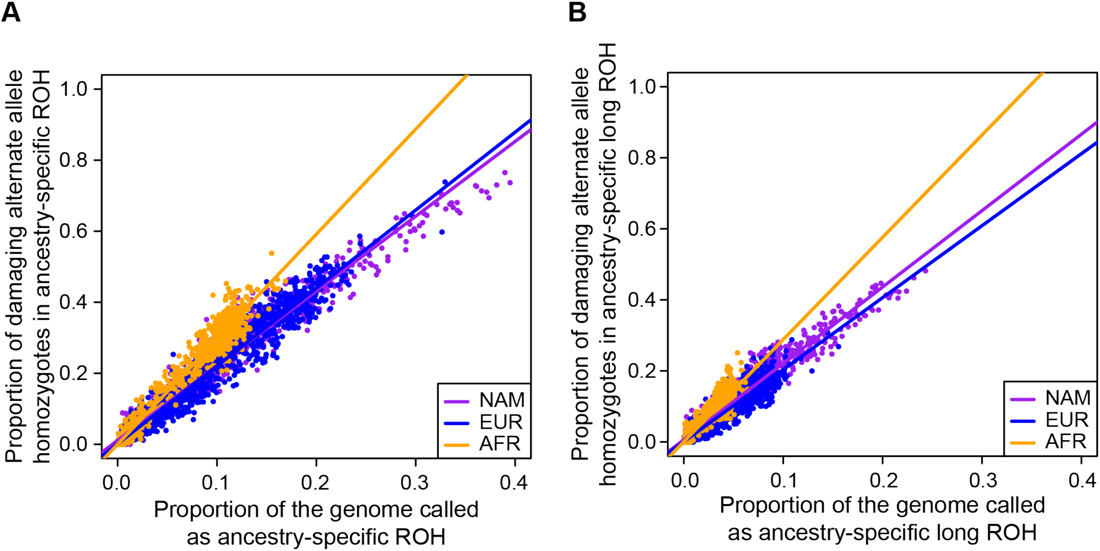
Deleterious homozygotes in ROH classes compared across ancestry. A direct comparison of the proportion of damaging homozygotes falling in ROH comprised of different ancestral haplotypes for (A) all ROH and (B) long ROH. EUR – European, colored blue; AFR – African, colored orange; and NAM – Native American, colored purple.

Consistent with our expectations, when analyzing all ROH (Fig 6A) we find a significant negative *β*_4_ = −0.763 (*p* < 2 × 10^−16^) and *β*_5_ = −0.852 (*p* < 2 × 10^−16^), indicating that the gain rate of damaging homozygotes in ROH on African ancestral haplotypes outpaces that of ROH on the other ancestral haplotypes. This pattern continues when considering only long ROH (*β*_4_ = −0.852, *p* < 2 × 10^−16^; *β*_5_ = −0.727, *p* < 2 × 10^−16^; Fig 6B) and smaller ROH (Table 4 and S2 Fig).

To check the robustness of these results, we reran these analyses using several other deleterious classification methods including SIFT [104, 105], Provean [106], and GERP [107]. Since GERP scores sites and not mutations, we restricted the GERP analysis to loci where the ancestral and derived states were inferred to high confidence. As this ancestral polarization results in discarding a large number of loci with ambiguous ancestral allele state, we also reran these analyses for Polyphen 2 [103], SIFT [104, 105], and Provean [106] restricted only to loci for which we have ancestral/derived state information. S3 Fig plots the inferred *β*_3_ for each of these analyses for each ROH size class and demonstrates qualitatively similar patterns as shown above.

We further re-analyzed a subset of the ROH and deleteriousness calls from Pemberton and Szpiech [74], which contains data on six admixed populations from the 1000 Genomes Project [108] and used CADD [109] scores as a deleteriousness prediction (S1 Text). After extracting the data relating to the admixed individuals from Pemberton and Szpiech [74] and calling local ancestries, we again find qualitatively similar patterns as above (S4 Fig).

Finally, since Pemberton and Szpiech [74] showed that these enrichment patterns appear to be driven by an abundance of homozygotes in ROH comprised of low-frequency alleles, we reanalyzed our data using categories of minor allele frequency (MAF) instead of deleteriousness. In order to determine MAF category, we use frequencies computed from all TOPMed Freeze 3 whole-genome sequencing data sets (dbGaP accession numbers phs000920, phs000921, phs001062, phs001032, phs000997, phs000993, phs001189, phs001211, phs001040, phs001024, phs000974, phs000956, phs000951, phs000946, phs000988, phs000964, phs000972, phs000954, and phs001143) forming a total sample size of *n =* 18,581. Using these allele frequencies, we categorize each polymorphic locus in a gene region (exons plus introns) into one of two categories: common (*MAF* ≥ 0.05) and rare (*MAF* < 0.05). We then fit the same models as above, except that instead of comparing the proportion of deleterious alternate allele homozygotes to benign homozygotes as a function of ROH coverage, we compare the number of minor allele homozygotes in the rare class to the common class.

We summarize the results of these analyses for each ancestral background, each ROH size class, and each low-frequency class in Fig 7. We find that ROH on African haplotype backgrounds are gain more low-frequency minor allele homozygotes per unit increase of ROH (and especially long class C ROH) compared to common minor allele homozygotes. Since low frequency alleles are enriched for deleterious variants relative to high frequency alleles, this result accords with our previous analyses.

**Fig 7.**
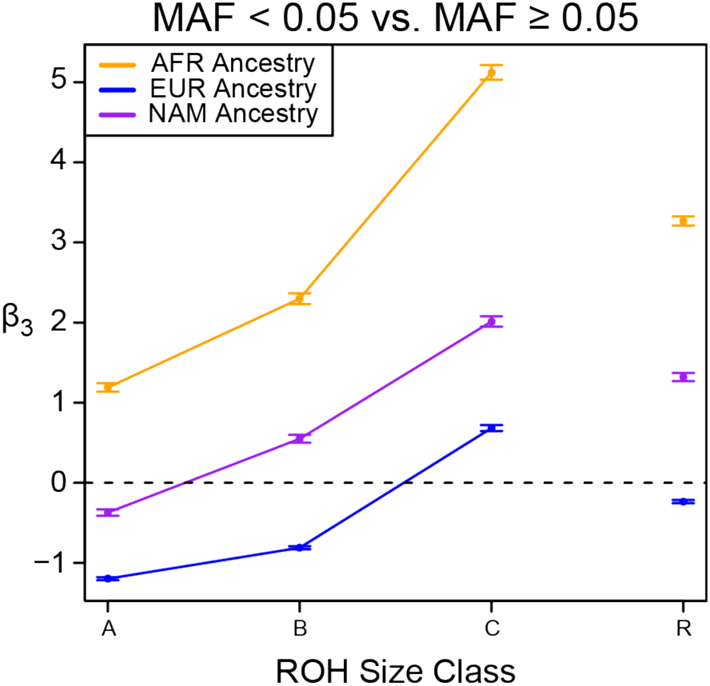
Enrichment of low-frequency variants across ROH sizes. The difference in rate of gain of low-frequency minor allele homozygotes (*MAF* < 0.05) compared to common minor allele homozygotes (*MAF* ≥ 0.05; *β*_3_ from regression analysis). ROH size classes: A – short, B – medium, C – long, R – all sizes. EUR – European, colored blue; AFR – African, colored orange; and NAM – Native American, colored purple. Error bars represent standard error of the regression coefficient.

## Discussion

The distribution of runs of homozygosity in individual genomes has provided insights into evolutionary, population, and medical genetics [1]. By examining their genomic location and prevalence in a population, we can learn about the history and adaptation of natural populations [2–39], and we can make discoveries about the genetic basis of complex phenotypes [40–66]. Given the importance of demographic history and socio-cultural practices in the generation of ROH in individual genomes, and their relationship to complex phenotypes including many genetic diseases, it naturally follows to study the distribution of deleterious alleles and their relationship to ROH.

Previous work has described the effect of demographic history on the distribution of deleterious alleles [98–101, 110, 111], including a few specifically investigating their relationship with runs of homozygosity [21, 38, 73, 74, 112, 113]. However, little work has been done on the relationship between deleterious alleles and ROH in admixed populations (although see [113]). Since there is evidence of very recent bottlenecks (which generate ROH) within admixed populations living in the Americas [88, 113], the relationship between ROH and the accumulation of deleterious homozygotes may provide valuable insights into the genetic basis of complex phenotypes in these individuals.

Here we analyzed 1,441 individuals across three admixed populations: African American, Puerto Rican, and Mexican American. We found that, consistent with other studies, the proportion of deleterious homozygotes found in ROH increases faster than the proportion of benign homozygotes as a function of total genomic ROH (Fig 4). However, we also proposed that ancestral haplotypes from populations with high deleterious *heterozygosity* would exhibit even greater increases of deleterious homozygotes per unit ROH. We reason that, under random mating, the larger number of low-frequency deleterious alleles in the population would largely segregate as heterozygotes, whereas, when a harsh bottleneck or consanguinity occurs, these mutations get paired IBD as homozygotes, concentrating more deleterious homozygotes within ROH. Indeed, we found that the genome-wide proportion of deleterious homozygotes in ROH on African ancestral haplotypes increased faster per unit ROH than on ether European or Native American ancestral haplotypes (Figs 5 and 6). These patterns are also consistent with population-specific worldwide patterns of deleterious homozygotes in ROH [74], where three of the five African populations analyzed had among the highest rates of enrichment in long ROH.

Whereas ROH on any haplotype background are associated with an increased rate of deleterious homozygotes, ROH on African haplotypes tend to have a larger share of the genome-wide deleterious homozygotes. Indeed, this accords with recent work that has independently associated increased ROH [65] and increased African ancestry [114] with reduced lung function. This suggests that these ROH on African haplotypes may play a particularly important role in the genetic architecture of complex phenotypes in admixed individuals, especially for populations with African ancestry that have undergone very harsh bottlenecks in the recent past.

## Methods

### Calling Local Ancestry

We used 90 African (YRI) individuals and 90 European (CEU) individuals for ancestry references (genotypes obtained from the Axiom^®^ Genotype Data Set at https://www.thermofisher.com/us/en/home/life-science/microarray-analysis/microarray-data-analysis/microarray-analysis-sample-data/axiom-genotype-data-set.html) and SNPs with less than 95% call rate were removed. For Native American reference genotypes we used 71 Native American individuals previously genotyped on the Axiom^®^ Genome-Wide LAT 1 array [115].

We then subset our 1,441 whole-genome sequences corresponding to sites found on the Axiom^®^ Genome-Wide LAT 1 array, leaving 765,321 markers. We then merge these data with our European (CEU), African (YRI), and Native American (NAM) reference panels, which overlapped at 434,145 markers. After filtering multi-allelic SNPs and SNPs with > 10% missing data, we obtained a final merged dataset of 428,644 markers. We phased this combined data set using SHAPEIT2 [116] and called local ancestry tracts jointly with RFMix [117] under a three-way admixture model based on the African, European, and Native American reference genotypes described above.

### Calling Runs of Homozygosity

We called runs of homozygosity using the program GARLIC v1.1.4 [102], which implements the ROH calling pipeline of [7], for each population separately on the full whole-genome call set, filtering only monomorphic sites. For the 475 African American (AA) individuals, this left 39,517,679 segregating sites; for the 483 Puerto Rican (PR) individuals, this left 31,961,900 segregating sites; and for the 483 Mexican American (MX) individuals, this left 30,744,389 segregating sites. Instead of asserting a single constant genotyping error rate (as in [7]), we used genotype quality scores provided with the WGS data to give GARLIC a per-genotype estimation of error. Using GARLIC’s rule of thumb parameter estimation, we chose analysis window sizes of 290 SNPs, 250 SNPs, and 210 SNPs for the AA, PR, and MX populations, respectively. Using GARLIC’s rule of thumb parameter estimation, we chose overlap fractions of 0.3688, 0.3553, and 0. 3528 for the AA, PR, and MX populations, respectively. GARLIC chose LOD score cutoffs of −47.5169, −70.1977, and −60.9221 for the AA, PR, and MX populations, respectively. Using a three-component Gaussian mixture model, GARLIC determined class A/B and class B/C size boundaries as 38,389 bps and 142,925 bps for AA; as 50,618 bps and 230,079 bps for PR; and 46,979 bps and 217,054 bps for MX.

### Calling Deleterious Alleles

Using the Whole Genome Sequencing Annotation (WGSA) pipeline [118] to generate annotation data, we extracted PolyPhen 2 [103], SIFT [104, 105], Provean [106], and GERP [107] scores for deleteriousness, as well as ancestral allele state and synonymous annotations and for all mutations in coding regions.

PolyPhen 2 generates three deleteriousness categories: Probably Damaging, Possible Damaging, and Benign. If a mutation has more than one PolyPhen2 classification (e.g. Benign and Probably Damaging), it is reassigned to have only the most damaging category of the group. All mutations that have a PolyPhen 2 prediction or that are synonymous, are then pooled into two separate categories: “damaging” and “benign.” All Probably Damaging or Possibly Damaging mutations are pooled into the “damaging” category, and all Benign and synonymous mutations are pooled into the “benign” category.

SIFT generates two deleteriousness categories, Intolerant and Tolerant, which we relabel “damaging” and “benign.” If a mutation has more than one SIFT classification, it is reassigned to have only the most damaging category of the group.

Provean generates two deleteriousness categories, Deleterious and Neutral, which we relabel “damaging” and “benign.” If a mutation has more than one Provean classification, it is reassigned to have only the most damaging category of the group.

GERP generates a numerical score at a given locus where a higher score indicates more deleteriousness for a derived allele at that locus. Here we focus on derived alleles that are very likely to be deleterious and combine all derived mutations at sites with GERP >= 6 into the category “damaging.” We form our “benign” category with all derived mutations with GERP < = 2.

## SUPPORTING INFORMATION

**S1 Fig. Deleterious versus benign homozygotes by ancestry for medium and small ROH**. The proportion of damaging (red) and benign (blue) homozygotes falling in ROH comprised of different ancestral haplotypes and size classes: (A) medium NAM ROH, (B) medium EUR ROH, (C) medium AFR ROH, (D) short NAM ROH, (E) short EUR ROH, and (F) short AFR ROH. EUR – European, AFR – African, and NAM – Native American.

**S2 Fig. Comparison of deleterious homozygotes between ancestries for medium and small ROH**. A direct comparison of the proportion of damaging homozygotes falling in ROH comprised of different ancestral haplotypes for (A) medium ROH and (B) short ROH. EUR – European, colored blue; AFR – African, colored orange; and NAM – Native American, colored purple.

**S3 Fig. Regression coefficients for analyses with other deleteriousness classifications**. The difference in slopes (*β*_3_ coefficients) between deleterious and benign categories across ROH size classes from re-analyses of the data using different deleteriousness classification schemes. (A) SIFT, (B) Provean, (C) GERP, (D) SIFT only with derived alleles, (E) Provean only with derived alleles, (F) Polyphen 2 only with derived alleles. ROH size classes: A – short, B – medium, C – long, R – all sizes. eUr – European, colored blue; AFR – African, colored orange; and NAM – Native American, colored purple.

**S4 Fig. Replication of findings using 1000 genomes data**. The identical analysis from Fig 4A-C except using the six admixed populations from the 1000 Genomes Project and CADD scores. (A) Native American ancestry, (B) European Ancestry, (C) African Ancestry.

**S5 Fig. The distribution of polarized Polyphen2 deleterious alleles across populations**. The number of (A) deleterious heterozygotes, (B) deleterious homozygotes, and (C) total deleterious alleles per individual using polarized Polyphen2 classifications. AA – African American, PR – Puerto Rican, MX – Mexican American.

**S6 Fig. The distribution of SIFT deleterious alleles across populations**. The number of (A) deleterious heterozygotes, (B) deleterious homozygotes, and (C) total deleterious alleles per individual using SIFT classifications. AA – African American, PR – Puerto Rican, MX – Mexican American.

**S7 Fig. The distribution of polarized SIFT deleterious alleles across populations**. The number of (A) deleterious heterozygotes, (B) deleterious homozygotes, and (C) total deleterious alleles per individual using polarized SIFT classifications. AA – African American, PR – Puerto Rican, MX – Mexican American.

**S8 Fig. The distribution of Provean deleterious alleles across populations**. The number of (A) deleterious heterozygotes, (B) deleterious homozygotes, and (C) total deleterious alleles per individual using Provean classifications. AA – African American, PR – Puerto Rican, MX – Mexican American.

**S9 Fig. The distribution of polarized Provean deleterious alleles across populations**. The number of (A) deleterious heterozygotes, (B) deleterious homozygotes, and (C) total deleterious alleles per individual using polarized Provean classifications. AA – African American, PR – Puerto Rican, MX – Mexican American.

**S10 Fig. The distribution of GERP deleterious alleles across populations**. The number of (A) deleterious heterozygotes, (B) deleterious homozygotes, and (C) total deleterious alleles per individual using GERP classifications. AA – African American, PR – Puerto Rican, MX – Mexican American.

**S1 Text. Processing of 1000 genomes data**.

## References

1. Ceballos FC, Joshi PK, Clark DW, Ramsay M, Wilson JF. Runs of homozygosity: windows into population history and trait architecture. Nature Reviews Genetics. 2018.

2. Broman KW, Weber JL. Long homozygous chromosomal segments in reference families from the centre d’tude du polymorphisme humain. Am J Hum Genet. 1999;65(6):1493–500. Epub 1999/12/01. doi: 10.1086/302661. PubMed PMID: 10577902; PubMed Central PMCID: PMCPMC1288359.

3. Gibson J, Morton NE, Collins A. Extended tracts of homozygosity in outbred human populations. Hum Mol Genet. 2006;15(5):789–95. Epub 2006/01/27. doi: 10.1093/hmg/ddi493. PubMed PMID: 16436455.

4. McQuillan R, Leutenegger AL, Abdel-Rahman R, Franklin CS, Pericic M, Barac-Lauc L, et al. Runs of homozygosity in European populations. Am J Hum Genet. 2008;83(3):359–72. Epub 2008/09/02. doi: 10.1016/j.ajhg.2008.08.007. PubMed PMID: 18760389; PubMed Central PMCID: PMCPMC2556426.

5. Kirin M, McQuillan R, Franklin CS, Campbell H, McKeigue PM, Wilson JF. Genomic runs of homozygosity record population history and consanguinity. PLoS One. 2010;5(11):e13996. Epub 2010/11/19. doi: 10.1371/journal.pone.0013996. PubMed PMID: 21085596; PubMed Central PMCID: PMCPMC2981575.

6. Nothnagel M, Lu TT, Kayser M, Krawczak M. Genomic and geographic distribution of SNP-defined runs of homozygosity in Europeans. Hum Mol Genet. 2010;19(15):2927–35. Epub 2010/05/14. doi: 10.1093/hmg/ddql98. PubMed PMID: 20462934.

7. Pemberton TJ, Absher D, Feldman MW, Myers RM, Rosenberg NA, Li JZ. Genomic patterns of homozygosity in worldwide human populations. Am J Hum Genet. 2012;91(2):275–92. Epub 2012/08/14. doi: 10.1016/j.ajhg.2012.06.014. PubMed PMID: 22883143; PubMed Central PMCID: PMCPMC3415543.

8. Blant A, Kwong M, Szpiech ZA, Pemberton TJ. Weighted likelihood inference of genomic autozygosity patterns in dense genotype data. BMC Genomics. 2017;18(1):928. Epub 2017/12/02. doi: 10.1186/S12864-017-4312-3. PubMed PMID: 29191164; PubMed Central PMCID: PMCPMC5709839.

9. Ben Halim N, Nagara M, Regnault B, Hsouna S, Lasram K, Kefi R, et al. Estimation of Recent and Ancient Inbreeding in a Small Endogamous Tunisian Community Through Genomic Runs of Homozygosity. Annals of Human Genetics. 2015;79(6):402–17. doi: 10.1111/ahg.l2131. PubMed PMID: WOS:000365399000003.

10. Kardos M, Luikart G, Allendorf FW. Measuring individual inbreeding in the age of genomics: marker-based measures are better than pedigrees. Heredity (Edinb). 2015;115(1):63–72. Epub 2015/06/11. doi: 10.1038/hdy.2015.17. PubMed PMID: 26059970; PubMed Central PMCID: PMCPMC4815495.

11. Mastrangelo S, Tolone M, Di Gerlando R, Fontanesi L, Sardina MT, Portolano B. Genomic inbreeding estimation in small populations: evaluation of runs of homozygosity in three local dairy cattle breeds. Animal. 2016;10(5):746–54. doi: 10.1017/S1751731115002943. PubMed PMID: WOS:000377125600003.

12. Kang JTL, Goldberg A, Edge MD, Behar DM, Rosenberg NA. Consanguinity Rates Predict Long Runs of Homozygosity in Jewish Populations. Hum Hered. 2016;82(3–4):87–102. Epub 2017/09/15. doi: 10.1159/000478897. PubMed PMID: 28910803; PubMed Central PMCID: PMCPMC5698150.

13. Meyer M, Kircher M, Gansauge MT, Li H, Racimo F, Mallick S, et al. A High-Coverage Genome Sequence from an Archaic Denisovan Individual. Science. 2012;338(6104):222–6. doi: 10.1126/science.1224344. PubMed PMID: WOS:000309712300037.

14. Castellano S, Parra G, Sanchez-Quinto FA, Racimo F, Kuhlwilm M, Kircher M, et al. Patterns of coding variation in the complete exomes of three Neandertals. P Natl Acad Sci USA. 2014;111(18):6666–71. doi: 10.1073/pnas.l405138111. PubMed PMID: WOS:000335477300045.

15. Prufer K, Racimo F, Patterson N, Jay F, Sankararaman S, Sawyer S, et al. The complete genome sequence of a Neanderthal from the Altai Mountains. Nature. 2014;505(7481):43–+. doi: 10.1038/nature12886. PubMed PMID: WOS:000329163300020.

16. Gamba C, Jones ER, Teasdale MD, McLaughlin RL, Gonzalez-Fortes G, Mattiangeli V, et al. Genome flux and stasis in a five millennium transect of European prehistory. Nature Communications. 2014;5. doi: ARTN 5257 10.1038/ncomms6257. PubMed PMID: WOS:000343985400006.

17. Prado-Martinez J, Sudmant PH, Kidd JM, Li H, Kelley JL, Lorente-Galdos B, et al. Great ape genetic diversity and population history. Nature. 2013;499(7459):471–5. doi: 10.1038/nature12228. PubMed PMID: WOS:000322157900039.

18. Xue YL, Prado-Martinez J, Sudmant PH, Narasimhan V, Ayub Q, Szpak M, et al. Mountain gorilla genomes reveal the impact of long-term population decline and inbreeding. Science. 2015;348(6231):242–5. doi: 10.1126/science.aaa3952. PubMed PMID: WOS:000352613700048.

19. Palkopoulou E, Mallick S, Skoglund P, Enk J, Rohland N, Li H, et al. Complete Genomes Reveal Signatures of Demographic and Genetic Declines in the Woolly Mammoth. Current Biology. 2015;25(10):1395–400. doi: 10.1016/j.cub.2015.04.007. PubMed PMID: WOS:000354785900035.

20. Curik I, Ferencakovic M, Solkner J. Inbreeding and runs of homozygosity: A possible solution to an old problem. Livest Sci. 2014;166:26–34. doi: 10.1016/j.livsci.2014.05.034. PubMed PMID: WOS:000340994000005.

21. Zhang Q, Guldbrandtsen B, Bosse M, Lund MS, Sahana G. Runs of homozygosity and distribution of functional variants in the cattle genome. BMC Genomics. 2015;16:542. Epub 2015/07/23. doi: 10.1186/sl2864-015-1715-x. PubMed PMID: 26198692; PubMed Central PMCID: PMCPMC4508970.

22. Howard JT, Tiezzi F, Huang Y, Gray KA, Maltecca C. Characterization and management of long runs of homozygosity in parental nucleus lines and their associated crossbred progeny. Genet Sel Evol. 2016;48. doi: ARTN 91 10.1186/sl2711-016-0269-y. PubMed PMID: WOS:000388541600002.

23. Manunza A, Noce A, Serradilla JM, Goyache F, Martinez A, Capote J, et al. A genome-wide perspective about the diversity and demographic history of seven Spanish goat breeds. Genet Sel Evol. 2016;48. doi: ARTN 52 10.1186/S12711-016-0229-6. PubMed PMID: WOS:000381071000001.

24. Gurgul A, Szmatola T, Topolski P, Jasielczuk I, Zukowski K, Bugno-Poniewierska M. The use of runs of homozygosity for estimation of recent inbreeding in Holstein cattle. J Appl Genet. 2016;57(4):527–30. doi: 10.1007/s13353-016-0337-6. PubMed PMID: WOS:000385423900013.

25. Peripolli E, Stafuzza NB, Munari DP, Lima ALF, Irgang R, Machado MA, et al. Assessment of runs of homozygosity islands and estimates of genomic inbreeding in Gyr (Bos indicus) dairy cattle. Bmc Genomics. 2018;19. doi: ARTN 34 10.1186/S12864-017-4365-3. PubMed PMID: WOS:000419680000001.

26. Forutan M, Mahyari SA, Baes C, Melzer N, Schenkel FS, Sargolzaei M. Inbreeding and runs of homozygosity before and after genomic selection in North American Holstein cattle. Bmc Genomics. 2018;19. doi: ARTN 98 10.1186/s12864-018-4453-z. PubMed PMID: WOS:000423444300001.

27. Kardos M, Qvarnstrom A, Ellegren H. Inferring Individual Inbreeding and Demographic History from Segments of Identity by Descent in Ficedula Flycatcher Genome Sequences. Genetics. 2017;205(3):1319–34. doi: 10.1534/genetics.116.198861. PubMed PMID: WOS:000395807200022.

28. Bortoluzzi C, Crooijmans R, Bosse M, Hiemstra SJ, Groenen MAM, Megens HJ. The effects of recent changes in breeding preferences on maintaining traditional Dutch chicken genomic diversity. Heredity (Edinb). 2018. Epub 2018/03/29. doi: 10.1038/s41437-018-0072-3. PubMed PMID: 29588508.

29. Bertolini F, Gandolfi B, Kim ES, Haase B, Lyons LA, Rothschild MF. Evidence of selection signatures that shape the Persian cat breed. Mamm Genome. 2016;27(3–4):144–55. doi: 10.1007/s00335-016-9623-1. PubMed PMID: WOS:000373308200005.

30. Boyko AR, Quignon P, Li L, Schoenebeck JJ, Degenhardt JD, Lohmueller KE, et al. A Simple Genetic Architecture Underlies Morphological Variation in Dogs. Plos Biol. 2010;8(8). doi: ARTN e1000451 10.1371/journal.pbio. 1000451. PubMed PMID: WOS:000281464500009.

31. vonHoldt BM, Pollinger JP, Earl DA, Knowles JC, Boyko AR, Parker H, et al. A genome-wide perspective on the evolutionary history of enigmatic wolf-like canids. Genome Research. 2011;21(8):1294–305. doi: 10.1101/gr.ll6301.110. PubMed PMID: WOS:000293335700009.

32. Pilot M, Dabrowski MJ, Hayrapetyan V, Yavruyan EG, Kopaliani N, Tsingarska E, et al. Genetic Variability of the Grey Wolf Canis lupus in the Caucasus in Comparison with Europe and the Middle East: Distinct or Intermediary Population? Plos One. 2014;9(4). doi: ARTN e93828 10.1371/journal.pone.0093828. PubMed PMID: WOS:000334160900054.

33. Friedenberg SG, Meurs KM, Mackay TFC. Evaluation of artificial selection in Standard Poodles using whole-genome sequencing. Mamm Genome. 2016;27(11–12):599–609. doi: 10.1007/s00335-016-9660-9. PubMed PMID: WOS:000388682800007.

34. Metzger J, Pfahler S, Distl O. Variant detection and runs of homozygosity in next generation sequencing data elucidate the genetic background of Lundehund syndrome. Bmc Genomics. 2016; 17. doi: ARTN 535 10.1186/S12864-016-2844-6. PubMed PMID: WOS:000381222500003.

35. Pedersen NC, Pooch AS, Liu H. A genetic assessment of the English bulldog. Canine genetics and epidemiology. 2016;3(1):6.

36. Dreger DL, Davis BW, Cocco R, Sechi S, Di Cerbo A, Parker HG, et al. Commonalities in Development of Pure Breeds and Population Isolates Revealed in the Genome of the Sardinian Fonni’s Dog. Genetics. 2016;204(2):737–55. doi: 10.1534/genetics.ll6.192427. PubMed PMID: WOS:000385871400028.

37. Wiener P, Sanchez-Molano E, Clements DN, Woolliams JA, Haskell MJ, Blott SC. Genomic data illuminates demography, genetic structure and selection of a popular dog breed. Bmc Genomics. 2017;18. doi: ARTN 609 10.1186/S12864-017-3933-x. PubMed PMID: WOS:000408036300001.

38. Sams AJ, Boyko AR. Fine-scale resolution and analysis of runs of homozygosity in domestic dogs. bioRxiv. 2018. doi: 10.1101/315770.

39. Kardos M, Akesson M, Fountain T, Flagstad O, Liberg O, Olason P, et al. Genomic consequences of intensive inbreeding in an isolated wolf population. Nat Ecol Evol. 2018;2(1):124–31. Epub 2017/11/22. doi: 10.1038/s41559-017-0375-4. PubMed PMID: 29158554.

40. Johnson EC, Evans LM, Keller MC. Relationships between estimated autozygosity and complex traits in the UK Biobank. bioRxiv. 2018. doi: 10.1101/291872.

41. Sheridan E, Wright J, Small N, Corry PC, Oddie S, Whibley C, et al. Risk factors for congenital anomaly in a multiethnic birth cohort: an analysis of the Born in Bradford study. Lancet. 2013;382(9901):1350–9. Epub 2013/07/09. doi: 10.1016/S0140-6736(13)61132-0. PubMed PMID: 23830354.

42. Bittles AH. Consanguineous marriages and congenital anomalies. Lancet. 2013;382(9901):1316–7. Epub 2013/07/09. doi: 10.1016/S0140-6736(13)61503-2. PubMed PMID: 23830356.

43. Scott EM, Halees A, Itan Y, Spencer EG, He Y, Azab MA, et al. Characterization of Greater Middle Eastern genetic variation for enhanced disease gene discovery. Nat Genet. 2016;48(9):1071–6. Epub 2016/07/19. doi: 10.1038/ng.3592. PubMed PMID: 27428751; PubMed Central PMCID: PMCPMC5019950.

44. Shami SA, Qaisar R, Bittles AH. Consanguinity and adult morbidity in Pakistan. Lancet. 1991;338(8772):954. Epub 1991/10/12. PubMed PMID: 1681304.

45. Puzyrev VP, Lemza SV, Nazarenko LP, Panphilov VI. Influence of genetic and demographic factors on etiology and pathogenesis of chronic disease in north Siberian aborigines. Arctic Med Res. 1992;51(3):136–42. Epub 1992/07/01. PubMed PMID: 1503580.

46. Ismail J, Jafar TH, Jafary FH, White F, Faruqui AM, Chaturvedi N. Risk factors for non-fatal myocardial infarction in young South Asian adults. Heart. 2004;90(3):259–63. Epub 2004/02/18. PubMed PMID: 14966040; PubMed Central PMCID: PMCPMC1768096.

47. Christofidou P, Nelson CP, Nikpay M, Qu L, Li M, Loley C, et al. Runs of Homozygosity: Association with Coronary Artery Disease and Gene Expression in Monocytes and Macrophages. Am J Hum Genet. 2015;97(2):228–37. Epub 2015/07/15. doi: 10.1016/j.ajhg.2015.06.001. PubMed PMID: 26166477; PubMed Central PMCID: PMCPMC4573243.

48. Simpson JL, Martin AO, Elias S, Sarto GE, Dunn JK. Cancers of the breast and female genital system: search for recessive genetic factors through analysis of human isolate. Am J Obstet Gynecol. 1981;141(6):629–36. Epub 1981/11/15. PubMed PMID: 7315892.

49. Lebel RR, Gallagher WB. Wisconsin consanguinity studies. II: Familial adenocarcinomatosis. Am J Med Genet. 1989;33(1):1–6. Epub 1989/05/01. doi: 10.1002/ajmg.l320330102. PubMed PMID: 2750776.

50. Rudan I. Inbreeding and cancer incidence in human isolates. Hum Biol. 1999;71(2):173–87. Epub 1999/05/01. PubMed PMID: 10222641.

51. Bacolod MD, Schemmann GS, Wang S, Shattock R, Giardina SF, Zeng Z, et al. The signatures of autozygosity among patients with colorectal cancer. Cancer Res. 2008;68(8):2610–21. Epub 2008/04/01. doi: 10.1158/0008-5472.CAN-07-5250. PubMed PMID: 18375840; PubMed Central PMCID: PMCPMC4383032.

52. Ujvari B, Klaassen M, Raven N, Russell T, Vittecoq M, Hamede R, et al. Genetic diversity, inbreeding and cancer. Proc Biol Sci. 2018;285(1875). Epub 2018/03/23. doi: 10.1098/rspb.2017.2589. PubMed PMID: 29563261; PubMed Central PMCID: PMCPMC5897632.

53. Krieger H. Inbreeding effects on metrical traits in Northeastern Brazil. Am J Hum Genet. 1969;21(6):537–46. Epub 1969/11/01. PubMed PMID: 5365755; PubMed Central PMCID: PMCPMC1706491.

54. Hurwich BJ, Nubani N. Blood pressures in a highly inbred community--Abu Ghosh, Israel. 1. Original survey. Isr J Med Sci. 1978;14(9):962–9. Epub 1978/09/01. PubMed PMID: 721424.

55. Saleh EA, Mahfouz AA, Tayel KY, Naguib MK, Bin-al-Shaikh NM. Hypertension and its determinants among primary-school children in Kuwait: an epidemiological study. East Mediterr Health J. 2000;6(2–3):333–7. Epub 2001/09/15. PubMed PMID: 11556020.

56. Rudan I, Smolej-Narancic N, Campbell H, Carothers A, Wright A, Janicijevic B, et al. Inbreeding and the genetic complexity of human hypertension. Genetics. 2003;163(3):1011–21. Epub 2003/03/29. PubMed PMID: 12663539; PubMed Central PMCID: PMCPMC1462484.

57. Campbell H, Carothers AD, Rudan I, Hayward C, Biloglav Z, Barac L, et al. Effects of genome-wide heterozygosity on a range of biomedically relevant human quantitative traits. Hum Mol Genet. 2007;16(2):233–41. Epub 2007/01/16. doi: 10.1093/hmg/ddl473. PubMed PMID: 17220173.

58. Keller MC, Miller G. Resolving the paradox of common, harmful, heritable mental disorders: which evolutionary genetic models work best? Behav Brain Sci. 2006;29(4):385–404; discussion 5-52. Epub 2006/11/11. doi: 10.1017/S0140525X06009095. PubMed PMID: 17094843.

59. Keller MC, Simonson MA, Ripke S, Neale BM, Gejman PV, Howrigan DP, et al. Runs of homozygosity implicate autozygosity as a schizophrenia risk factor. PLoS Genet. 2012;8(4):e1002656. Epub 2012/04/19. doi: 10.1371/journal.pgen.1002656. PubMed PMID: 22511889; PubMed Central PMCID: PMCPMC3325203.

60. Gandin I, Faletra F, Faletra F, Carella M, Pecile V, Ferrero GB, et al. Excess of runs of homozygosity is associated with severe cognitive impairment in intellectual disability. Genet Med. 2015;17(5):396–9. Epub 2014/09/19. doi: 10.1038/gim.2014.118. PubMed PMID: 25232855.

61. Mukherjee S, Guha S, Ikeda M, Iwata N, Malhotra AK, Pe’er I, et al. Excess of homozygosity in the major histocompatibility complex in schizophrenia. Hum Mol Genet. 2014;23(22):6088–95. Epub 2014/06/20. doi: 10.1093/hmg/ddu308. PubMed PMID: 24943592; PubMed Central PMCID: PMCPMC4204767.

62. Iourov IY, Vorsanova SG, Korostelev SA, Zelenova MA, Yurov YB. Long contiguous stretches of homozygosity spanning shortly the imprinted loci are associated with intellectual disability, autism and/or epilepsy. Mol Cytogenet. 2015;8:77. Epub 2015/10/20. doi: 10.1186/sl3039-015-0182-z. PubMed PMID: 26478745; PubMed Central PMCID: PMCPMC4608298.

63. Ghani M, Reitz C, Cheng R, Vardarajan BN, Jun G, Sato C, et al. Association of Long Runs of Homozygosity With Alzheimer Disease Among African American Individuals. JAMA Neurol. 2015;72(11):1313–23. Epub 2015/09/15. doi: 10.1001/jamaneurol.2015.1700. PubMed PMID: 26366463; PubMed Central PMCID: PMCPMC4641052.

64. McQuillan R, Eklund N, Pirastu N, Kuningas M, McEvoy BP, Esko T, et al. Evidence of inbreeding depression on human height. PLoS Genet. 2012;8(7):e1002655. Epub 2012/07/26. doi: 10.1371/journal.pgen. 1002655. PubMed PMID: 22829771; PubMed Central PMCID: PMCPMC3400549.

65. Joshi PK, Esko T, Mattsson H, Eklund N, Gandin I, Nutile T, et al. Directional dominance on stature and cognition in diverse human populations. Nature. 2015;523(7561):459–62. Epub 2015/07/02. doi: 10.1038/nature14618. PubMed PMID: 26131930; PubMed Central PMCID: PMCPMC4516141.

66. Lyons EJ, Frodsham AJ, Zhang L, Hill AV, Amos W. Consanguinity and susceptibility to infectious diseases in humans. Biol Lett. 2009;5(4):574–6. Epub 2009/03/28. doi: 10.1098/rsbl.2009.0133. PubMed PMID: 19324620; PubMed Central PMCID: PMCPMC2684220.

67. Pritchard JK. Are rare variants responsible for susceptibility to complex diseases? Am J Hum Genet. 2001;69(1):124–37. Epub 2001/06/19. doi: 10.1086/321272. PubMed PMID: 11404818; PubMed Central PMCID: PMCPMC1226027.

68. Pritchard JK, Cox NJ. The allelic architecture of human disease genes: common disease-common variant…or not? Hum Mol Genet. 2002;11(20):2417–23. Epub 2002/09/28. PubMed PMID: 12351577.

69. Carlson CS, Eberle MA, Kruglyak L, Nickerson DA. Mapping complex disease loci in whole-genome association studies. Nature. 2004;429(6990):446–52. Epub 2004/05/28. doi: 10.1038/nature02623. PubMed PMID: 15164069.

70. Freimer N, Sabatti C. The use of pedigree, sib-pair and association studies of common diseases for genetic mapping and epidemiology. Nat Genet. 2004;36(10):1045–51. Epub 2004/09/30. doi: 10.1038/ngl433. PubMed PMID: 15454942.

71. Boyle EA, Li YI, Pritchard JK. An Expanded View of Complex Traits: From Polygenic to Omnigenic. Cell. 2017;169(7):1177–86. Epub 2017/06/18. doi: 10.1016/j.cell.2017.05.038. PubMed PMID: 28622505; PubMed Central PMCID: PMCPMC5536862.

72. Reich DE, Lander ES. On the allelic spectrum of human disease. Trends Genet. 2001;17(9):502–10. Epub 2001/08/30. PubMed PMID: 11525833.

73. Szpiech ZA, Xu J, Pemberton TJ, Peng W, Zollner S, Rosenberg NA, et al. Long runs of homozygosity are enriched for deleterious variation. Am J Hum Genet. 2013;93(1):90–102. Epub 2013/06/12. doi: 10.1016/j.ajhg.2013.05.003. PubMed PMID: 23746547; PubMed Central PMCID: PMCPMC3710769.

74. Pemberton TJ, Szpiech ZA. Relationship between Deleterious Variation, Genomic Autozygosity, and Disease Risk: Insights from The 1000 Genomes Project. Am J Hum Genet. 2018;102(4):658–75. Epub 2018/03/20. doi: 10.1016/j.ajhg.2018.02.013. PubMed PMID: 29551419.

75. Colby SL, Ortman JM. Projections of the size and composition of the US population: 2014 to 2060: Population estimates and projections. 2017.

76. Popejoy AB, Fullerton SM. Genomics is failing on diversity. Nature. 2016;538(7624):161–4. doi: DOI 10.1038/538161a. PubMed PMID: WOS:000386671000016.

77. Martin AR, Gignoux CR, Walters RK, Wojcik GL, Neale BM, Gravel S, et al. Human Demographic History Impacts Genetic Risk Prediction across Diverse Populations. Am J Hum Genet. 2017;100(4):635–49. Epub 2017/04/04. doi: 10.1016/j.ajhg.2017.03.004. PubMed PMID: 28366442; PubMed Central PMCID: PMCPMC5384097.

78. Verdu P, Austerlitz F, Estoup A, Vitalis R, Georges M, Thery S, et al. Origins and genetic diversity of pygmy hunter-gatherers from Western Central Africa. Curr Biol. 2009;19(4):312–8. Epub 2009/02/10. doi: 10.1016/j.cub.2008.12.049. PubMed PMID: 19200724.

79. Verdu P, Rosenberg NA. A general mechanistic model for admixture histories of hybrid populations. Genetics. 2011;189(4):1413–26. Epub 2011/10/05. doi: 10.1534/genetics.111.132787. PubMed PMID: 21968194; PubMed Central PMCID: PMCPMC3241432.

80. Via M, Gignoux CR, Roth LA, Fejerman L, Galanter J, Choudhry S, et al. History shaped the geographic distribution of genomic admixture on the island of Puerto Rico. PLoS One. 2011;6(1):e16513. Epub 2011/02/10. doi: 10.1371/journal.pone.0016513. PubMed PMID: 21304981; PubMed Central PMCID: PMCPMC3031579.

81. Gravel S. Population genetics models of local ancestry. Genetics. 2012;191(2):607–19. Epub 2012/04/12. doi: 10.1534/genetics. 112.139808. PubMed PMID: 22491189; PubMed Central PMCID: PMCPMC33 74321.

82. Moreno-Estrada A, Gravel S, Zakharia F, McCauley JL, Byrnes JK, Gignoux CR, et al. Reconstructing the population genetic history of the Caribbean. PLoS Genet. 2013;9(11):e1003925. Epub 2013/11/19. doi: 10.1371/journal.pgen.l003925. PubMed PMID: 24244192; PubMed Central PMCID: PMCPMC3828151 Ancestry.com, 23andMe’s “Roots into the Future” project, and Personalis, Inc. He is on the medical advisory board of Invitae and Med-tek. None of these entities played any role in the project or research results reported here.

83. Gravel S, Zakharia F, Moreno-Estrada A, Byrnes JK, Muzzio M, Rodriguez-Flores JL, et al. Reconstructing Native American migrations from whole-genome and whole-exome data. PLoS Genet. 2013;9(12):e1004023. Epub 2014/01/05. doi: 10.1371/journal.pgen.l004023. PubMed PMID: 24385924; PubMed Central PMCID: PMCPMC3873240.

84. Goldberg A, Verdu P, Rosenberg NA. Autosomal admixture levels are informative about sex bias in admixed populations. Genetics. 2014;198(3):1209–29. Epub 2014/09/07. doi: 10.1534/genetics.114.166793. PubMed PMID: 25194159; PubMed Central PMCID: PMCPMC4224161.

85. Verdu P, Pemberton TJ, Laurent R, Kemp BM, Gonzalez-Oliver A, Gorodezky C, et al. Patterns of admixture and population structure in native populations of Northwest North America. PLoS Genet. 2014;10(8):e1004530. Epub 2014/08/15. doi: 10.1371/journal.pgen.l004530. PubMed PMID: 25122539; PubMed Central PMCID: PMCPMC4133047.

86. Homburger JR, Moreno-Estrada A, Gignoux CR, Nelson D, Sanchez E, Ortiz-Tello P, et al. Genomic Insights into the Ancestry and Demographic History of South America. PLoS Genet. 2015;11(12):e1005602. Epub 2015/12/05. doi: 10.1371/journal.pgen.l005602. PubMed PMID: 26636962; PubMed Central PMCID: PMCPMC4670080.

87. Baharian S, Barakatt M, Gignoux CR, Shringarpure S, Errington J, Blot WJ, et al. The Great Migration and African-American Genomic Diversity. PLoS Genet. 2016;12(5):e1006059. Epub 2016/05/28. doi: 10.1371/journal.pgen.l006059. PubMed PMID: 27232753; PubMed Central PMCID: PMCPMC4883799.

88. Browning SR, Browning BL, Daviglus ML, Durazo-Arvizu RA, Schneiderman N, Kaplan RC, et al. Ancestry-specific recent effective population size in the Americas. PLoS Genet. 2018;14(5):e1007385. Epub 2018/05/26. doi: 10.1371/journal.pgen.l007385. PubMed PMID: 29795556.

89. Zhu X, Tang H, Risch N. Admixture mapping and the role of population structure for localizing disease genes. Adv Genet. 2008;60:547–69. Epub 2008/03/25. doi: 10.1016/S0065-2660(07)00419-1. PubMed PMID: 18358332.

90. Basu A, Tang H, Arnett D, Gu CC, Mosley T, Kardia S, et al. Admixture mapping of quantitative trait loci for BMI in African Americans: evidence for loci on chromosomes 3q, 5q, and 15q. Obesity (Silver Spring). 2009;17(6):1226–31. Epub 2009/07/09. doi: 10.1038/oby.2009.24. PubMed PMID: 19584881; PubMed Central PMCID: PMCPMC2929755.

91. Cheng CY, Kao WH, Patterson N, Tandon A, Haiman CA, Harris TB, et al. Admixture mapping of 15,280 African Americans identifies obesity susceptibility loci on chromosomes 5 and X. PLoS Genet. 2009;5(5):e1000490. Epub 2009/05/23. doi: 10.1371/journal.pgen.l000490. PubMed PMID: 19461885; PubMed Central PMCID: PMCPMC2679192.

92. Cheng CY, Reich D, Coresh J, Boerwinkle E, Patterson N, Li M, et al. Admixture mapping of obesity-related traits in African Americans: the Atherosclerosis Risk in Communities (ARIC) Study. Obesity (Silver Spring). 2010;18(3):563–72. Epub 2009/08/22. doi: 10.1038/oby.2009.282. PubMed PMID: 19696751; PubMed Central PMCID: PMCPMC2866099.

93. Torgerson DG, Gignoux CR, Galanter JM, Drake KA, Roth LA, Eng C, et al. Case-control admixture mapping in Latino populations enriches for known asthma-associated genes. Journal of Allergy and Clinical Immunology. 2012;130(1):76–82. e12.

94. Shriner D. Overview of admixture mapping. Curr Protoc Hum Genet. 2013;Chapter 1:Unit 123. Epub 2013/01/15. doi: 10.1002/0471142905.hg0123s76. PubMed PMID: 23315925; PubMed Central PMCID: PMCPMC3556814.

95. Galanter JM, Gignoux CR, Torgerson DG, Roth LA, Eng C, Oh SS, et al. Genome-wide association study and admixture mapping identify different asthma-associated loci in Latinos: the Genes-environments & Admixture in Latino Americans study. J Allergy Clin Immunol. 2014;134(2):295–305. Epub 2014/01/11. doi: 10.1016/j.jaci.2013.08.055. PubMed PMID: 24406073; PubMed Central PMCID: PMCPMC4085159.

96. Spear ML, Hu D, Pino-Yanes M, Huntsman S, Eng C, Levin AM, et al. A Genome-wide Association and Admixture Mapping Study of Bronchodilator Drug Response in African Americans with Asthma. bioRxiv. 2017. doi: 10.1101/157198.

97. Mak AC, White MJ, Eckalbar WL, Szpiech ZA, Oh SS, Pino-Yanes M, et al. Whole Genome Sequencing of Pharmacogenetic Drug Response in Racially Diverse Children with Asthma. Am J Respir Crit Care Med. 2018. Epub 2018/03/07. doi: 10.1164/rccm.201712-25290C. PubMed PMID: 29509491.

98. Lohmueller KE, Indap AR, Schmidt S, Boyko AR, Hernandez RD, Hubisz MJ, et al. Proportionally more deleterious genetic variation in European than in African populations. Nature. 2008;451(7181):994–7. Epub 2008/02/22. doi: 10.1038/nature06611. PubMed PMID: 18288194; PubMed Central PMCID: PMCPMC2923434.

99. Tennessen JA, Bigham AW, O’Connor TD, Fu W, Kenny EE, Gravel S, et al. Evolution and functional impact of rare coding variation from deep sequencing of human exomes. Science. 2012;337(6090):64–9. Epub 2012/05/19. doi: 10.1126/science.l219240. PubMed PMID: 22604720; PubMed Central PMCID: PMCPMC3708544.

100. Fu W, O’Connor TD, Jun G, Kang HM, Abecasis G, Leal SM, et al. Analysis of 6,515 exomes reveals the recent origin of most human protein-coding variants. Nature. 2013;493(7431):216–20. Epub 2012/12/04. doi: 10.1038/nature11690. PubMed PMID: 23201682; PubMed Central PMCID: PMCPMC3676746.

101. Henn BM, Botigue LR, Bustamante CD, Clark AG, Gravel S. Estimating the mutation load in human genomes. Nat Rev Genet. 2015;16(6):333–43. Epub 2015/05/13. doi: 10.1038/nrg3931. PubMed PMID: 25963372; PubMed Central PMCID: PMCPMC4959039.

102. Szpiech ZA, Blant A, Pemberton TJ. GARLIC: Genomic Autozygosity Regions Likelihood-based Inference and Classification. Bioinformatics. 2017;33(13):2059–62. Epub 2017/02/17. doi: 10.1093/bioinformatics/btxl02. PubMed PMID: 28205676; PubMed Central PMCID: PMCPMC5870576.

103. Adzhubei IA, Schmidt S, Peshkin L, Ramensky VE, Gerasimova A, Bork P, et al. A method and server for predicting damaging missense mutations. Nat Methods. 2010;7(4):248–9. Epub 2010/04/01. doi: 10.1038/nmeth0410-248. PubMed PMID: 20354512; PubMed Central PMCID: PMCPMC2855889.

104. Ng PC, Henikoff S. SIFT: Predicting amino acid changes that affect protein function. Nucleic Acids Res. 2003;31(13):3812–4. Epub 2003/06/26. PubMed PMID: 12824425; PubMed Central PMCID: PMCPMC168916.

105. Kumar P, Henikoff S, Ng PC. Predicting the effects of coding non-synonymous variants on protein function using the SIFT algorithm. Nat Protoc. 2009;4(7):1073–81. Epub 2009/06/30. doi: 10.1038/nprot.2009.86. PubMed PMID: 19561590.

106. Choi Y, Chan AP. PROVEAN web server: a tool to predict the functional effect of amino acid substitutions and indels. Bioinformatics. 2015;31(16):2745–7. Epub 2015/04/09. doi: 10.1093/bioinformatics/btvl95. PubMed PMID: 25851949; PubMed Central PMCID: PMCPMC4528627.

107. Cooper GM, Stone EA, Asimenos G, Program NCS, Green ED, Batzoglou S, et al. Distribution and intensity of constraint in mammalian genomic sequence. Genome Res. 2005;15(7):901–13. Epub 2005/06/21. doi: 10.1101/gr.3577405. PubMed PMID: 15965027; PubMed Central PMCID: PMCPMC1172034.

108. Genomes Project C, Auton A, Brooks LD, Durbin RM, Garrison EP, Kang HM, et al. A global reference for human genetic variation. Nature. 2015;526(7571):68–74. Epub 2015/10/04. doi: 10.1038/nature15393. PubMed PMID: 26432245; PubMed Central PMCID: PMCPMC4750478.

109. Kircher M, Witten DM, Jain P, O’Roak BJ, Cooper GM, Shendure J. A general framework for estimating the relative pathogenicity of human genetic variants. Nat Genet. 2014;46(3):310–5. Epub 2014/02/04. doi: 10.1038/ng.2892. PubMed PMID: 24487276; PubMed Central PMCID: PMCPMC3992975.

110. Lohmueller KE. The distribution of deleterious genetic variation in human populations. Curr Opin Genet Dev. 2014;29:139–46. Epub 2014/12/03. doi: 10.1016/j.gde.2014.09.005. PubMed PMID: 25461617.

111. Pedersen CT, Lohmueller KE, Grarup N, Bjerregaard P, Hansen T, Siegismund HR, et al. The Effect of an Extreme and Prolonged Population Bottleneck on Patterns of Deleterious Variation: Insights from the Greenlandic Inuit. Genetics. 2017;205(2):787–801. Epub 2016/12/03. doi: 10.1534/genetics.116.193821. PubMed PMID: 27903613; PubMed Central PMCID: PMCPMC5289852.

112. Mezzavilla M, Vozzi D, Badii R, Alkowari MK, Abdulhadi K, Girotto G, et al. Increased rate of deleterious variants in long runs of homozygosity of an inbred population from Qatar. Hum Hered. 2015;79(1):14–9. Epub 2015/02/28. doi: 10.1159/000371387. PubMed PMID: 25720536.

113. Mooney J, Huber C, Service S, Sul JH, Marsden C, Zhang Z, et al. Understanding the Hidden Complexity of Latin American Population Isolates. bioRxiv. 2018. doi: 10.1101/340158.

114. Pino-Yanes M, Thakur N, Gignoux CR, Galanter JM, Roth LA, Eng C, et al. Genetic ancestry influences asthma susceptibility and lung function among Latinos. J Allergy Clin Immunol. 2015;135(1):228–35. Epub 2014/10/11. doi: 10.1016/j.jaci.2014.07.053. PubMed PMID: 25301036; PubMed Central PMCID: PMCPMC4289103.

115. Drake KA, Torgerson DG, Gignoux CR, Galanter JM, Roth LA, Huntsman S, et al. A genome-wide association study of bronchodilator response in Latinos implicates rare variants. J Allergy Clin Immunol. 2014;133(2):370–8. Epub 2013/09/03. doi: 10.1016/j.jaci.2013.06.043. PubMed PMID: 23992748; PubMed Central PMCID: PMCPMC3938989.

116. Delaneau O, Zagury JF, Marchini J. Improved whole-chromosome phasing for disease and population genetic studies. Nat Methods. 2013;10(1):5–6. Epub 2012/12/28. doi: 10.1038/nmeth.2307. PubMed PMID: 23269371.

117. Maples BK, Gravel S, Kenny EE, Bustamante CD. RFMix: a discriminative modeling approach for rapid and robust local-ancestry inference. Am J Hum Genet. 2013;93(2):278–88. Epub 2013/08/06. doi: 10.1016/j.ajhg.2013.06.020. PubMed PMID: 23910464; PubMed Central PMCID: PMCPMC3738819.

118. Liu X, White S, Peng B, Johnson AD, Brody JA, Li AH, et al. WGSA: an annotation pipeline for human genome sequencing studies. J Med Genet. 2016;53(2):111–2. Epub 2015/09/24. doi: 10.1136/jmedgenet-2015-103423. PubMed PMID: 26395054; PubMed Central PMCID: PMCPMC5124490.

